# A generator of morphological clones for plant species

**DOI:** 10.1101/108530

**Authors:** Ilya Potapov, Marko Järvenpää, Markku Åkerblom, Pasi Raumonen, Mikko Kaasalainen

## Abstract

Detailed and realistic tree form generators have numerous applications in ecology and forestry. Here, we present an algorithm for generating morphological tree “clones” based on the detailed reconstruction of the laser scanning data, statistical measure of similarity, and a plant growth algorithm with simple stochastic rules. The algorithm is designed to produce tree forms, i.e. morphological clones, similar as a whole (coarse-grain scale), but varying in minute details of organization (fine-grain scale). We present a general procedure for obtaining these morphological clones. Although we opted for certain choices in our algorithm, its various parts may vary depending on the application. Namely, we have shown that specific multi-purpose procedural stochastic growth model can be algorithmically adjusted to produce the morphological clones replicated from the target experimentally measured tree. For this, we have developed a statistical measure of similarity (structural distance) between any given pair of trees, which allows for the comprehensive comparing of the tree morphologies in question by means of empirical distributions describing geometrical and topological features of a tree. Our algorithm can be used in variety of applications and contexts for exploration of the morphological potential of the growth models, arising in all sectors of plant science research.

**Summary Statement:** We present an algorithmic framework, based on the Bayesian inference, for generating morphological tree clones using a combination of stochastic growth models and experimentally derived tree structures.

## I. Introduction

Models for plant architecture attract significant attention due to their ability to assist the empirical studies in ecology, plant biology, forestry, and agronomy (Prusinkiewicz, 2004). The modeling activity is especially useful in research since it arises as fruitful collaboration between specialists in different fields of studies: computer scientists, mathematicians, and biologists (Fourcaud et al., 2008).

Modeling plant architecture is approached from many directions. Some progress has been achieved in synthesis of realistic plant forms in the field of computer graphics (Palubicki et al., 2009; Pirk et al., 2012; Stava et al., 2014). These models, although based on heuristic rules of growth, produce realistic shape outcomes in a fast and efficient manner, which is usually dictated by the application of this approach, that is natural sceneries in computer visualization. Heuristic growth rules of the procedural models for graphics applications are not firmly based on biological principles, but nevertheless elucidate some algorithmic properties of the growth process (for example, recursive (Hallé et al., 1978) vs. self-organizing (Sachs and Novoplansky, 1995; Palubicki et al., 2009) character of architecture development).

However, the most promising plant architectural models are so called functional-structural plant models (FSPM), also known as “virtual plants” (Room et al., 1996; Sievänen et al., 2000; Godin et al., 2004), because this type of models allows for a balanced description between morphological and functional/physiological properties of a plant. Thus, it is capable of connecting the external abiotic factors (e.g. radiation, temperature and soil) and the most vital functions of a plant organism (such as photosynthesis, respiration, and water and salts uptake) with its structural characteristics (Prusinkiewicz, 2004; Fourcaud et al., 2008).

Nevertheless, biologically relevant architectural plant models rely on data in a form of empirically fitted functions and parameters that correspond to a particular species and/or certain site conditions (Mäkelä and Hari, 1986; Rauscher et al, 1990; Perttunen et al., 1996; Lacointe, 2000). Thus, the change in these conditions requires re-calibration of the models, which is done in a manual fashion every time the model is simulated for the new conditions. Strong dependence on data, where each simulation would be calibrated automatically by data, is limited by both computation time and lack of the fast measurement and processing systems allowing for a detailed 3D morphological reconstruction of the real plant/tree.

The most recent advances in laser scanning techniques allow for fast and non-destructive measurement of trees with subsequent reconstruction of various characteristics depending on application (e.g. (Rosell et al., 2009; Van Leeuwen and Nieuwenhuis, 2010)). Most of such studies dedicated to reconstruction of 3D point clouds obtained from laser scanning measurements deal with overall characteristics, such as height, width, and volume of stems/crowns, leaf index, biomass etc., resembling traditional destructive methods of measurement (Rosell et al., 2009; Rutzinger et al., 2010). However, the detailed precise geometrical and topological reconstruction with the preserved tree architecture as is, is rarely sought after.

In this work, we use a fast, precise, automatic, and comprehensive reconstruction algorithm initially presented in (Raumonen et al., 2013) and further developed and tested in (Calders et al., 2015). The algorithm reliably reconstructs a quantitative structure model (QSM), which contains all geometrical and topological characteristics of the object tree. Input for the method is the 3D point cloud, sufficiently covering the tree, obtained from the terrestrial laser scanning measurements (TLS) and no additional allometric relations used for estimation of the branch proportions (as in (Xu et al., 2007; Livny et al., 2010)) are needed. Compared to other similar techniques (e.g. (Xu et al., 2007; Livny et al., 2010; Preuksakarn et al., 2010)) this method requires few parameters and no user interaction and reconstructs the tree surface with subsequent cylinder (or any other geometrical primitive) approximation, which is usually consistent with theoretical plant growth models. The reconstruction algorithm has been validated in several studies with several different tree species and different scanner instruments (Calders et al., 2015; Hackenberg et al., 2015; Kaasalainen et al., 2014; Raumonen et al., 2015; Smith et al., 2014). There are other published QSM reconstruction methods from TLS data that can produce similar quality QSMs, at least (Hackenberg et al., 2015).

In this work, we utilize an inverse iterative procedure to optimize model’s parameters as to match the (empirical) distribution of structural features of the simulated stochastic tree models (FSPM, graphical or other) to that of the tree reconstructed from the laser scanning data. Meanwhile, we formulate a measure of similarity of the tree structures grounded in tomographic analysis of the structural distributions (e.g. Radon transform) (Kaasalainen, 2008; Bracewell, 1990). Finally, the optimal parameter set produces morphological “clone” trees with similar overall structure, but varying minute details of organization.

Recently, we have reported a proof-of-concept study where we used reconstruction of a pine tree and the corresponding FSPM (named LIGNUM (Perttunen et al., 1996; Sievanen et al., 2008)) to demonstrate the practical feasibility of the approach (Potapov et al., 2016). In this work, however, we develop a unifying interface for our procedure and use general-purpose fast procedural tree growth model from (Palubicki et al., 2009), since such a simple procedural model is easier to adapt (it is simple, fast, and efficient) for technical experimentation with the whole algorithm. Additionally, similar algorithmic pipeline was reported in (Stava et al, 2014) for procedural tree growth models in the context of graphics synthesis. However, in our approach we see the tree growth as a random process and, consequently, apply corresponding statistical methods for measuring the similarity between trees. Moreover, in our algorithm the special concern is on biologically relevant description, hence, the careful choice of the reconstruction algorithm; possibility to use FSPM to relate physiological parameters to the morphogenetic processes in trees; and no extra structures improving visual properties of trees but not supported by empirical observation (e.g. leaves).

## II. Results

### Algorithm overview

Our approach is based upon five distinct parts:

1. *Quantitative Structure Model (QSM)* is a reconstruction of a tree model from 3D point clouds obtained from terrestrial laser scanning measurements (TLS). Here we use specific algorithm for such reconstruction reported in (Raumonen et al., 2013) and (Calders et al., 2015) but others could be used as well.
2. *Stochastic Structure Model (SSM)* is a tree growth model that is chosen depending on the application. There are no limitations on the class of the model, except it must produce measurable 3D branching structure.
3. *Structural data set (U)* is a collection of structural features (empirical distributions) to be compared between QSM and SSM. Importantly, *U* data sets must be determined in the same way both for QSM and SSM.
4. *Measure of structural dissimilarity, or structural distance D*_*S*_, is a measure of discrepancy between any two data sets, in other words, *D*_*S*_(*U*_*1*_, *U*_*2*_) results in a value quantifying how much different the two data sets *U*_*1*_ and *U*_*2*_ are.
5. *Optimization algorithm* is a numerical routine capable of finding a minimum of any given function by varying its arguments (Newton algorithm, genetic algorithm, simulated annealing etc.)

The connection between these components is outlined in Fig. 1 with explanation in the text below.

**Figure 1:**
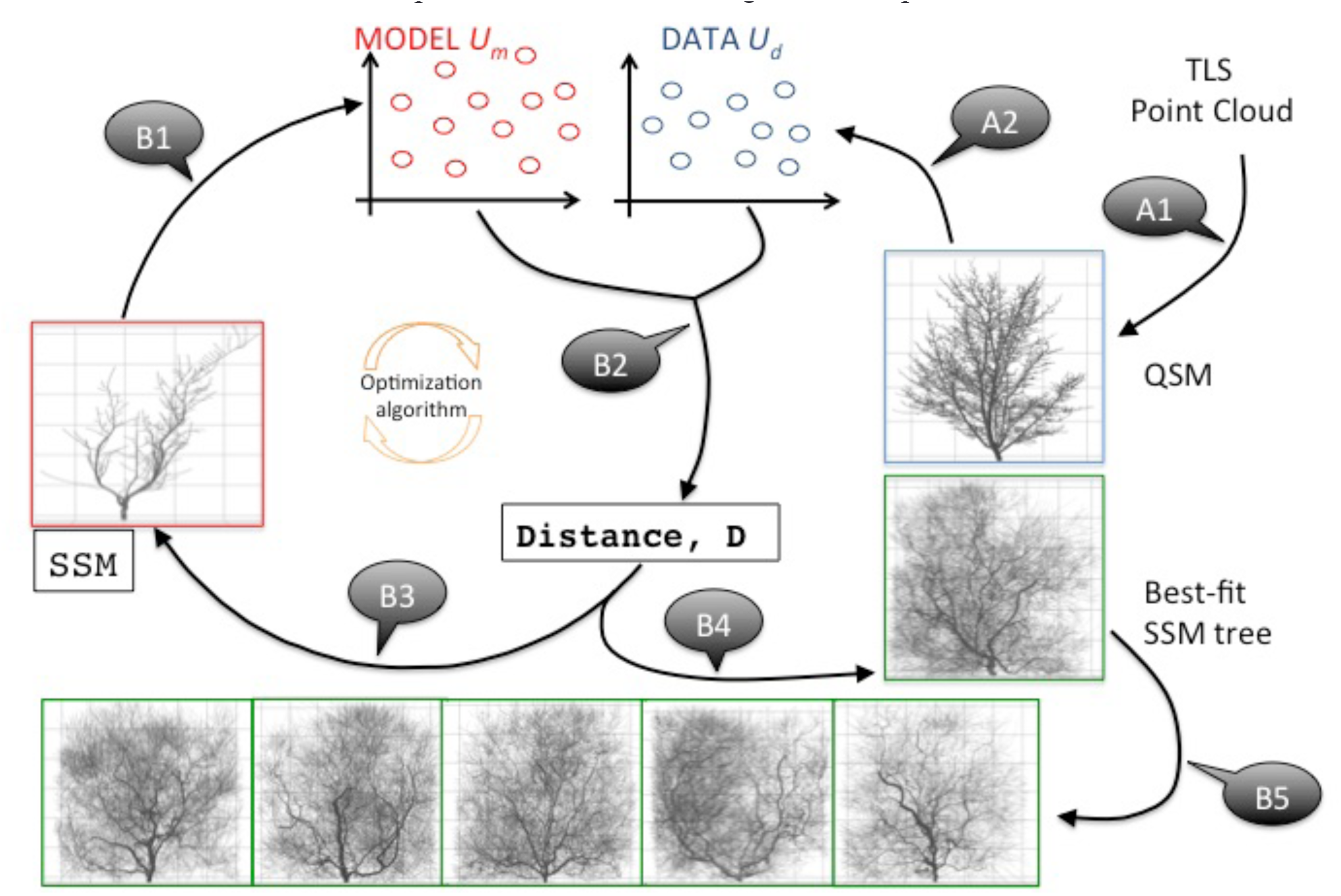
The algorithm outline (see explanation in the text).

The algorithm outline (Fig. 1):

Preparation stage A:

A1: build QSM from TLS.

A2: extract *U*_*d*_ from QSM.

Main cycle B:

B1: simulate SSM for the fixed parameters and extract *U*_*m*_.

B2: compare *U*_*m*_ and *U*_*d*_ getting an estimation of the distance *D* between them.

B3: change SSM parameters trying to decrease *D*, go to B1 or stop and go to B4 (changing of the parameters and stopping criteria depend on any particular realization of the optimization routine).

B4: simulate SSM with the “best-fit” parameter values corresponding to the smallest found *D*.

B5: loose the randomness of the best-fit SSM and generate morphological clones.

At the preparation stage, the QSM is formed from the TLS point cloud (A1). The detailed description of this process is reported in (Raumonen et al., 2013; Calders et al., 2015). The resultant QSM contains all geometrical and topological features needed to form the empirical distributions *U*_*d*_. The distributions can be formed for several tree individuals if they are close by shape to ensure the sample size.

At the main cycle of the algorithm, the empirical distribution *U*_*m*_ is formed from the simulated SSM tree (B1). Next, *U*_*m*_ is compared against *U*_*d*_ using the measure of distance (B2). The optimization routine iteratively minimizes the distance value every time changing the parameter values of SSM (B3), simulating SSM, and repeating the cycle from B1. After the stopping criteria of the optimization routine (number of iterations, minimal allowed distance etc.) are met, the algorithm stops and produces the best-fit SSM tree (B4). The best-fit SSM with different random sequences produces different outcomes – morphological clones.

In Materials and methods, we describe each of the main components of the algorithm in further detail.

### Preliminary observations

In the beginning of our analysis, we make several important notes about the target QSM structure. The shrub-like shape of this reconstruction model produces several major branches emanating from the initial part of the trunk connected to the ground. All these branches can be equally assigned with the order *w* = 0 (continuation of the trunk; see the definition of the topological order *w* in Materials and methods), however, the heuristic algorithm of the tree reconstruction from the TLS data (Raumonen et al., 2013) at every branching point chooses the thickest pathway to determine the actual trunk (it is roughly the thickest pathway, although the actual algorithm specifies much more complicated rules, see (Raumonen et al., 2013) for details). This has the following implications.

First, tree with the zero and first order branches has a skewed shape (Fig. 2A), since only one of the trunk candidate branches becomes the actual trunk (*w* = 0) whereas the rest of them become the first order branches (*w* = 1). The asymmetry of the form appears due to the branches attached to the actual trunk and assigned with *w* = 1, because other similarly scaled and attached to other trunk candidates branches become effectively the branches of order *w* = 2. Second, due to the aforementioned asymmetry the data sets for the first order branches have a modular structure: large scaled trunk-like branches along with the smaller ones. Third, we observe that the overall shape of the subject QSM can be approximated by the branches of the topological orders *w* ≤ 2 as it can be seen from Fig. 1B. Namely, with orders *w =* 0 and 1 the shape of the tree seems to be underrepresented (mainly due to the shape asymmetry), while with orders *w =* 0, 1, 2, and 3 the smaller twigs just fill in the spatial gaps between the major branches. This makes the analysis and form fitting a more complex task as compared with the tree shapes resulting from the growth with strong apical dominance (e.g. pine trees; see (Potapov et al., 2016)).

**Figure 2:**
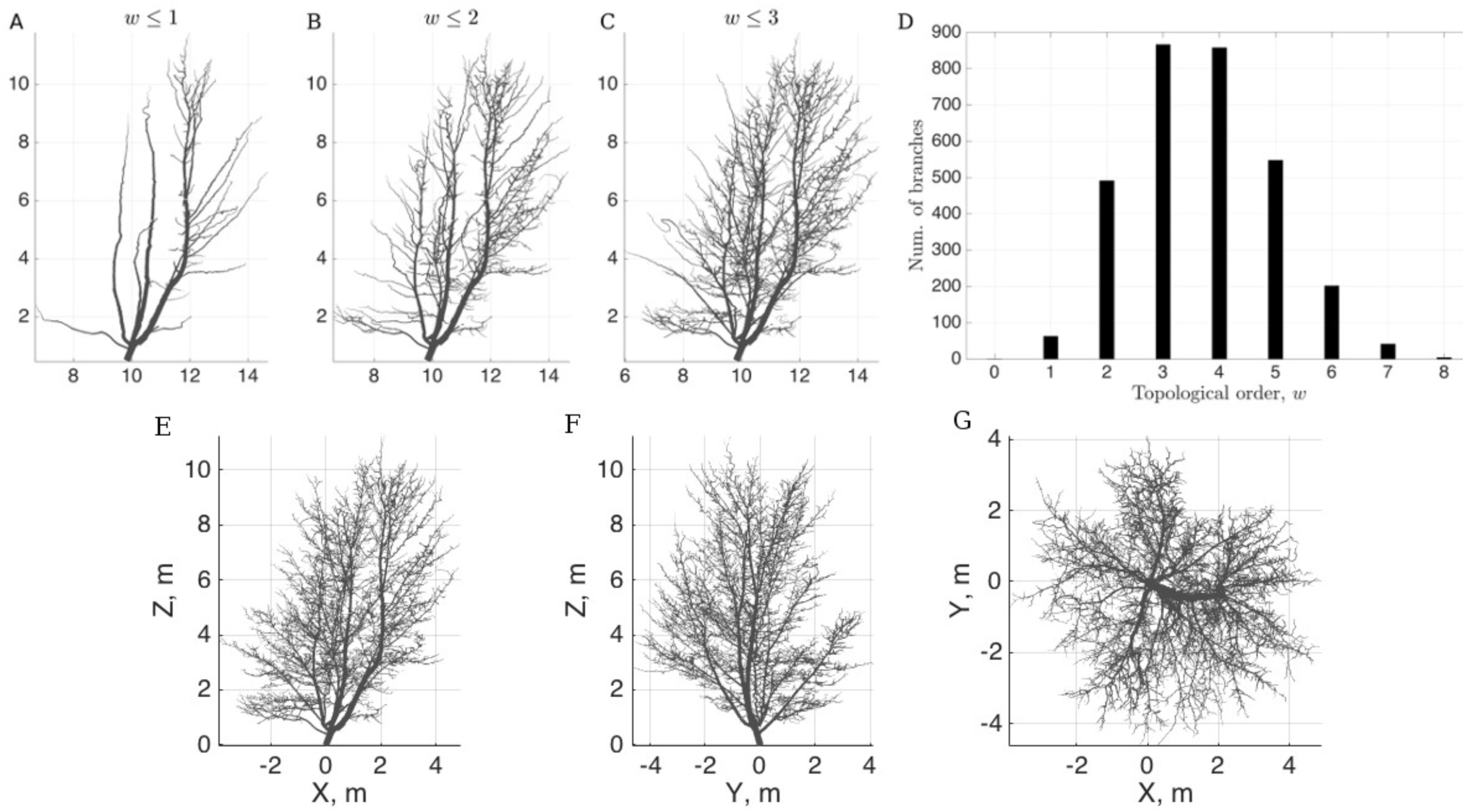
The target QSM structure. (A) *w* = 0, 1; (B) *w* = 0, 1, 2; (C) *w* = 0, 1, 2, 3; (D) distribution of the topological orders *w* of the QSM. Full QSM tree: XZ-projection (E), YZ-projection (F), and XY-projection (G).

Another point to consider is the underlying statistical properties. For example, it is impossible to draw any branch statistics from the single instance of the trunk (*w =* 0), while there are plenty of samples for the higher order branches. Given that the overall shape is mainly governed by the lower order branches, one must compromise between the main, shape determining branches with lower abundance and less important, but numerous, higher order branches (Fig. 2D).

Finally, the branch-related (*B*, see Materials and methods for the notation) data sets do not provide sufficient information for the width of the branches and their curvature in space. Moreover, although the *B* set has some information on the width (*R*_*f*_, *L*_*t*_), it is less abundant than the similar and more detailed information contained in the segment-related (*S*, see Materials and methods) data sets. However, the *B* data set has information on the structure of the emanating pattern of a branch, that is, the spatial location of its lateral buds/branching points (*L*_*a*_), and its angular properties, which, in turn, can be substituted with the biologically plausible growth algorithm.

Therefore, we begin our analysis with *S*^*0,1*^ data sets as *w* = 0, 1 branches represent the main structural frame of the tree: without its valid approximation the whole tree cannot be considered fitted.

### Basic values of the parameters

First, we run the optimization within each of the parameter groups *I – V* (see Materials and methods) to determine the basic values of the parameters. These basic values represent choices that generate a viable tree structure with proportions and scale approximately equal to those of the target QSM. Each optimization run takes the best parameters for the group optimized at the previous step. The target distributions *U* for these runs are *S*^*0,1*^. Note that this exercise serves a basic exploration of the model’s behavior, which can be (partially) replaced, for example, by the expert guesses for the parameter values or some calibration process (if the model is designed for specific purposes and/or species).

Second, based on these preliminary results we determine the most influential parameters for each of the group and combine them in a single optimization set up. Several independent optimization runs were taken in order to determine the most influential parameters. For example, we found that the angular properties vary the least among these runs, whereas the apical dominance requires subtler adjustments (as can be understood from the complex structure of the target QSM).

### Low order topological adjustment of the shape

After these initial manipulations, we obtained a model with 11 parameters and good fit of the trunk and first order branches (Fig. 3C; *d*_*h*_ = 0.05, *d*_*g*_ = 0.42, *d*_*c*_ = 0.57). However, the overall form of the resulting minimal score tree does not resemble the target QSM due to its rosette-shape (Fig. 3A, B). A closer look at the tree reveals that the higher order branches (*w* > 1) are mainly responsible for the formation of the rosette-shape of the tree, i.e. the orders which were not subject to the optimization (Fig. 3). This example demonstrates the contribution of the higher order branches to the overall tree shape, which suggests using the scatters of these orders in further optimization steps. Moreover, the branch-related features, such as the angular properties of branches of order *w* > 1, were not captured well (Fig. 3E), although similar order segment-related features show right stochastic tendencies (Fig. 3D) generated automatically by the growth algorithm of the SSM. This further stipulates usage of *B* scatters of orders *w* > 1.

**Figure 3:**
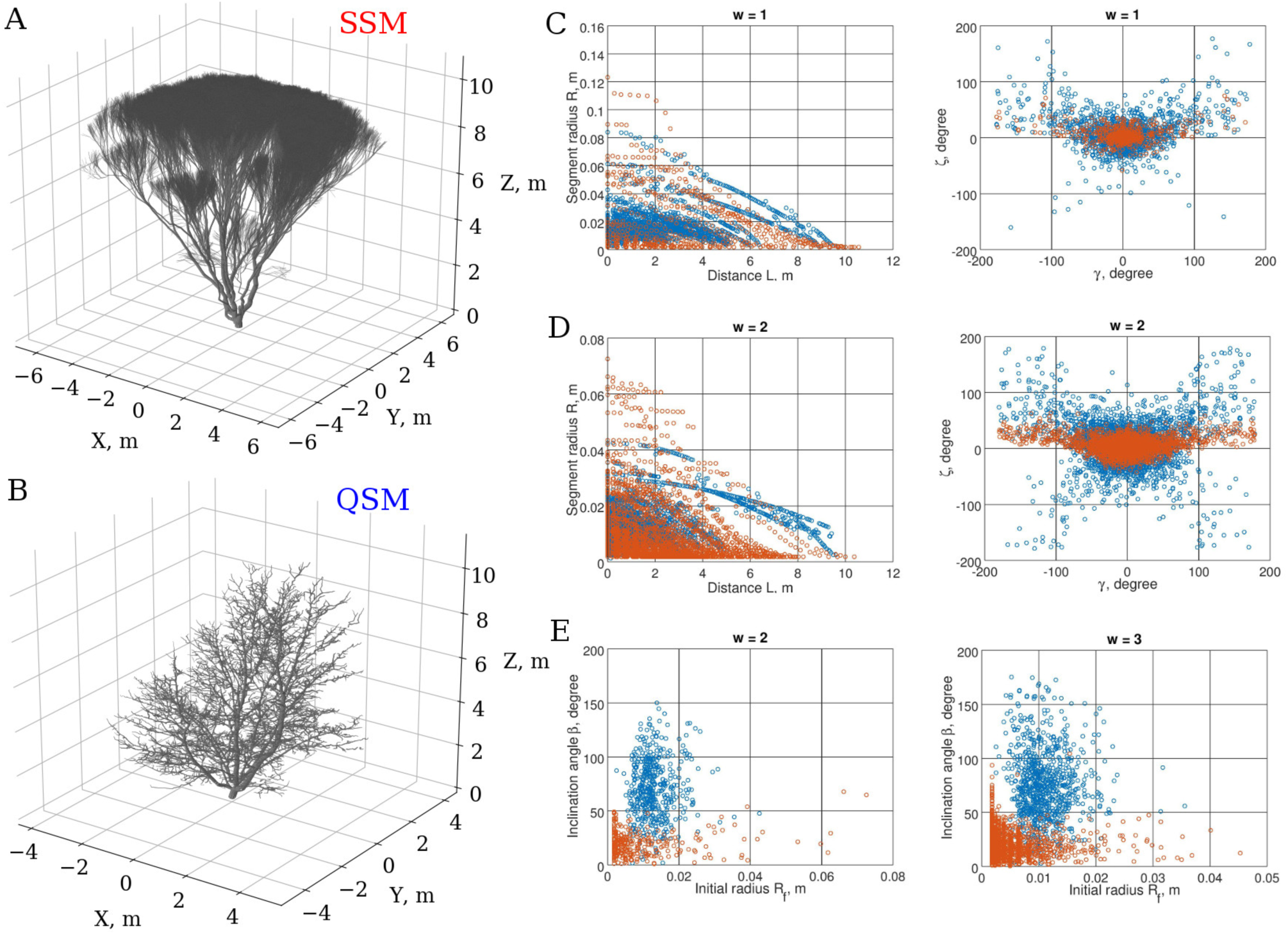
The rosette-shape SSM resulting from the adjustment of the low order (*S*^0,1^) scatters. (A) The SSM tree; (B) the target QSM; (C) some *S*^0,1^ scatters used in the optimization; (D) higher order (*w* = 2) *S*-scatters; (E) higher order (*w* = 2, 3) *B*-scatters. Note that the scatters in (D) and (E) were not used in the optimization. SSM/QSM scatters are shown in red/blue.

### Higher order topological adjustment

The increase in number of the structural feature tables is coupled with the increase in number of distinct distance values. Although the optimization of the mean distance value hinders the improvement for each target table, the low order as well as high order branches need to be fitted to the corresponding branches of the target QSM as we have shown above (Fig. 3). To reduce the number of distinct feature tables for the optimization we further utilize the merged data sets resulting in two joint *S* and *B* tables for all topological orders (see Materials and methods).

Thus, we opted for *S*^0,1^ and *B*^2,3,4^ merged data sets in the next run of optimization to account for the higher order branch variability (Fig. 4, *d*_*h*_ = 0.08, *d*_*g*_ = 0.20, *d*_*c*_ = 0.68). No longer we can see the rosette-shape due to the correct account of the angular properties of the higher order (*w* > 1) branches (Fig. 4E). The poor convergence of the branch linear dimensions (radii, lengths etc.) present in the branch-related tables might be due to the parameter choice of the model. Namely, the small proportion of branches demonstrating right *R*_*f*_ values (Fig. 4E) appears to be the result of the fixed segment length, we opted for as a compromise between reality and computational complexity (the QSM minimal segment length is close to zero, median is 0.06 m). Noteworthy is the similar span of the curvature data points of SSM and QSM for *w* = 1, 2 (Fig. C and D), although *w* = 2 branch curvature was not subject to the optimization. Additionally, due to the lack of the orientation landmark in the feature data sets our best-fit SSM is fitted to the target QSM with accuracy of the rotation around Z-axis (this could be adjusted, for example, by associating South direction with a coordinate axis).

**Figure 4:**
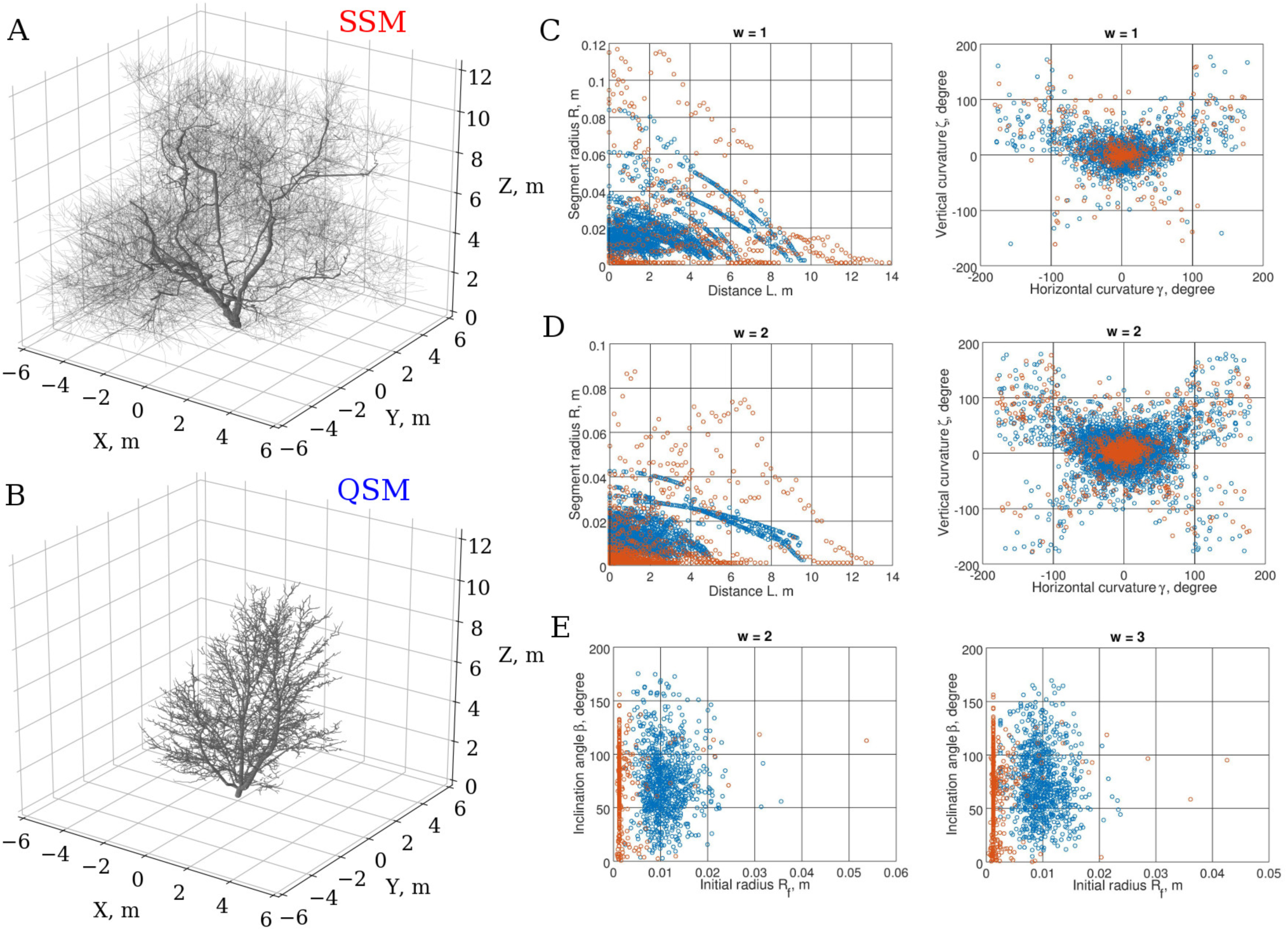
Low and high order adjustment of the stochastic feature tables. The best-fit SSM is obtained through optimization against *S*^0,1^ and *B*^2,3,4^ merged feature data sets. (A) The best-fit SSM tree, (B) the target QSM tree, (C) some projection scatters from *S*^1^, (D) *S*^2^ projection scatters, (E) *B*^2^ and *B*^3^ projection scatters.

### Clonal nature of the best-fit SSM

Due to the highly discrete and stochastic nature of the tree growth, the structural distance hyper-surface in the space of the parameters is extremely abrupt (Fig. 5A). Hence, finding the global minima of such surface is not a trivial task (the classical smooth function optimizers are not suitable in this case, while stochastic discrete optimizers, like the genetic algorithm, seem to be more appropriate). Moreover, the hyper-surface itself is a stochastic entity changing every time the new sample of random numbers is used for a particular SSM growth realization. Therefore, any best-fit SSM is the best for a particular realization of this stochastic process: one needs to study variability of the tree shape and the chances are that other SSM growth realization can produce a lower distance value (Fig. 5B). We call these many realizations of the SSM growth *morphological tree clones*.

**Figure 5:**
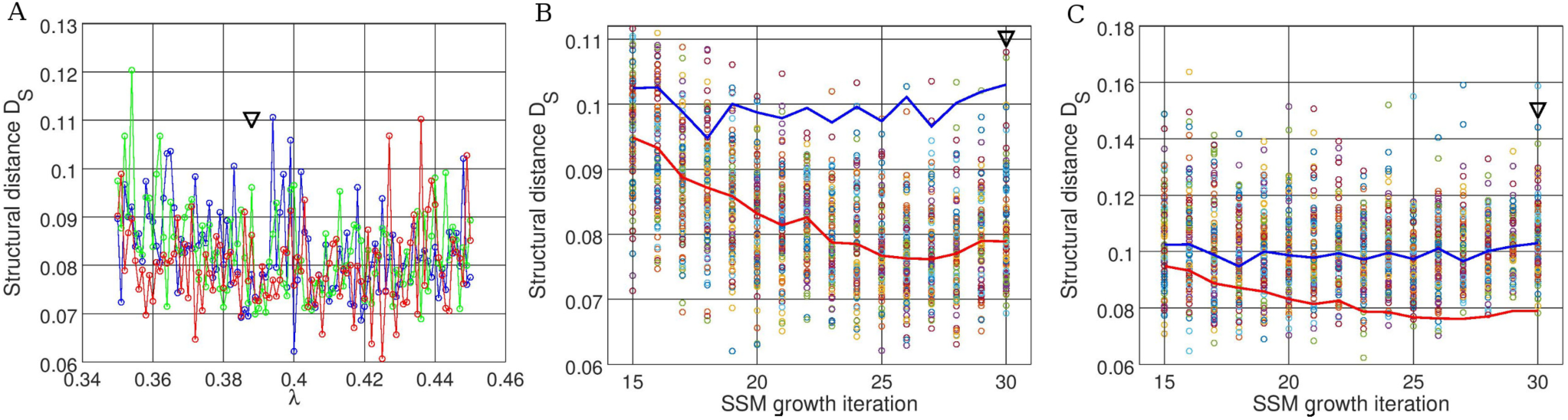
Stochastic structure distance profiles in the parameter space. (A) Three realizations of the distance hyper-surface projection along a dimensionless parameter λ of the SSM, controlling the apical dominance of a tree (the shown fragment of the projection with the step of 0.001 approximates 30% of the allowed variability of the parameter during optimization, which was [0.35, 0.65]). (B) Structural distance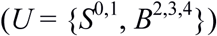 values for 100 randomly generated SSM trees for each value of a discrete SSM parameter, i.e. number of growth iterations (red line connects the median points of the distance distributions for each parameter value; blue line shows the same median distance profile but for the disturbed system from (C)). (C) Same as in (B), but *U* = *S*^0,1^ (blue line is the median profile; red line is from (B)). The SSM is the best-fit SSM from Fig. 4; the black arrow indicates the parameter value of the best-fit SSM.

The structural distance profile depends not only on the parameters of the SSM, but the choice of the structural data sets. For example, in Fig. 5B and C the median distance profile is depicted given *U* = 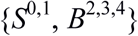 (red line) and *U* = *S*^0,1^ (blue line). In the given parameter span the latter seems to be more flattened and lifted compared to the former. The addition of the *B*^2,3,4^ data set might be seen as a perturbation to the distance profile changing the landscape properties (like minima). In our simulations we maintain the global parameter boundaries, which allows for the search within the full available space. However, we sequentially improve the model characteristics by perturbing the system, i.e. changing the parameters, their intervals, and the *U* data sets to address problematic parts of the SSM such that at every next optimization run the genetic algorithm is instructed to search around the previous best point using the initial ranges (see Materials and methods).

Given the considerations above about the nature of the structural distance hyper-surface, the further study of the morphological clones is needed. Specifically, the variability and plausibility of the clonal shapes need to be addressed. For example, the clones must be further selected as to produce realistic tree shapes (especially, when the general purpose SSM is used, like in this study), although we could not find any unrealistic trees out of the best-fit SSM in our analysis. Additionally, the variability of the clones is to be calibrated, for instance, by the analysis of the natural/QSM clonal individuals.

### Morphological tree clones

The quintessence of our work is the generation of the morphological clones. In our pipeline, this occupies the last stage (see Fig. 1, B5). After the optimization is finished and the best-fit SSM is found, one can further randomize the outcome of SSM by letting the random number generator produce different sequences every time SSM is run. As a result, the different realizations of SSM should constitute the morphological clone generator yielding structural copies close to QSM and to each other and *varying* in fine detail of organization of their branches. In other words, the coarse-grain structure is repeated in each clone (and possibly grasps that of the target QSM), whereas the fine-grain structure varies.

**Figure 6:**
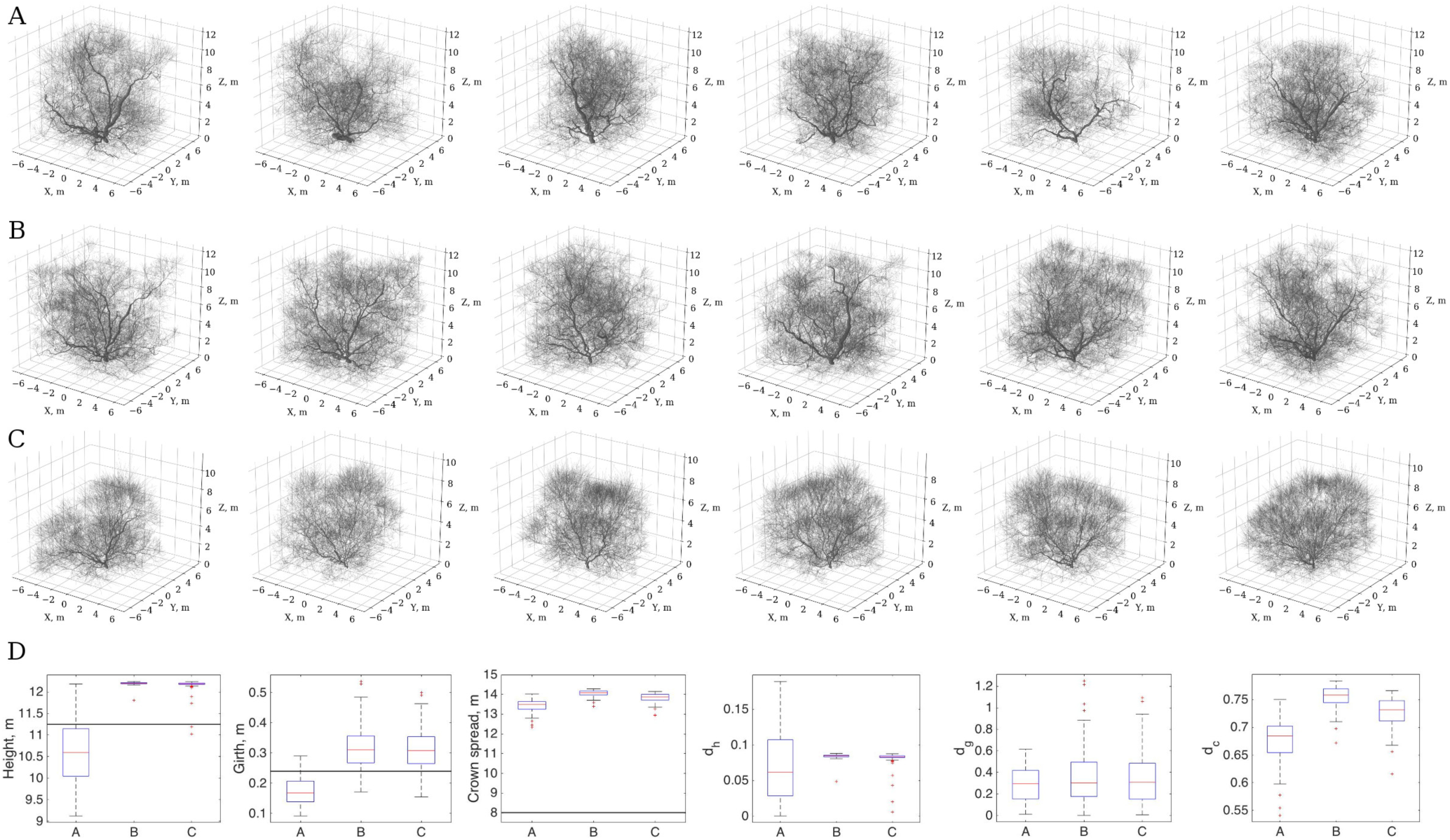
Morphological clones generated from the best-fit SSM. The best-fit SSM was found using the higher topological order adjustments (Fig. 4) with number of growth iterations 30 (A), 26 (B), and 18 (C). The height, girth, crown spread, and classical metrics distributions are shown in (D) for the clones in (A), (B), and (C) (the total number of generated clones for each case is *n* = 100). The black horizontal line indicates the corresponding measure of the target QSM.

We demonstrate visualization of six clones for three distinct cases in Fig. 6. One can see the fine-grain variation in the structure in each panel of the figure, although the overall (coarse-grain) structure is preserved and presumably captures that of the target maple QSM (Fig. 2). The three models are: the one found during the optimization process (Fig. 6A), the one minimizing the sample median distance profile for 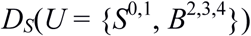 shown in Fig. 5B and one minimizing the sample median profile *D*_*S*_(*U* = *S*^0,1^) from Fig. 5C.

Out of 100 simulated clones for each case, we can see that the best-fit SSM obtained directly as the optimization outcome (Fig. 6A) produces larger proportion of individual trees exhibiting the three standard allometric measures closer to those of QSM (Fig. 6D). However, we argue that such simple description of a tree as using the allometric measures cannot be exhaustive enough to capture both the overall structure and its fine details.

The height statistics have the largest variability but by the visual inspection of the drawn clones in Fig. 6 one can see that this variability does not exert significant alterations of the Z axis span and the trees seem to have even heights. Perhaps, the way we calculate the height of a tree produces such large deviations in each particular case, which makes it a non-robust estimator.

Similarly, the girth estimation, although being captured decently, produces large errors *d*_*g*_, which seems to be a result of variation in its linear dimensions (Fig. 6D). The girth dimension spans a small proportion of the dimension of the whole tree: from several to tens of centimeters compared to meters of the whole tree. This makes the girth specific error look gigantic (exceeding in some cases 100%) and thus non-robust as well.

The crown spread measure shows significant variation (Fig. 6D). We believe that this takes place due to the environment of the real tree the QSM was reconstructed from, which was not modeled appropriately in the SSM. Namely, the environmental effects (positions relative to the sun, as the tree grows in the Northern country, animals, winds, neighboring trees etc.) might cause systematic influences exerted on the shape of the QSM tree. These influences were not accounted for in the SSM, which was allowed to grow in any direction, limited by the light conditions, existing branches of the same tree, and global boundaries of the available space. In addition to the environment influences, there are TLS measurement and QSM reconstruction errors, arising from the physical limitations of the instrumental technique and stochasticity of the QSM formation, respectively.

Finally, the true understanding of the variability of any measures of the morphological clones comes with the measurements of the real clones. Carrying out control experiments with QSM reconstructed from the real clonal individuals can only assess the variability. These real clone controlled experiments can further identify whether the obtained variability is large/small for the given species/clones and lead to the adjustment of the optimization parameters.

### Bayes-Forest toolbox

We have further developed a unified interface using Matlab facilitating exploration, drawing, optimization, and simulation of SSM and QSM as well as study of the morphological tree clones. Our interface allows for faster and easier manipulation of the required data, models, and optimization routines from the Matlab Optimization Toolbox, using only the required elements of otherwise complex Matlab configuration for the analysis.

The Bayes-Forest toolbox is freely available at http://math.tut.fi/inversegroup/app/bayesforest/v1/. We also encourage the plant and computer scientists’ community to expand their efforts using the toolbox with other species and models. Such a systematic approach can further be useful in tinkering the best options for creating QSM, SSM, and construction of the structural data sets.

## III. Discussion

In this work, we described an algorithmic pipeline aimed at producing stochastic structural replicas, or morphological “clones”, of trees from a QSM tree (data from TLS reconstruction) and a complimentary SSM tree (analytical/procedural growth model). The pipeline is based on an iterative minimization of a distance between morphological structures. The distance is based on construction of the structural data sets of the tree morphologies and subsequent measure of their discrepancy using the ideas of distribution tomography analysis. The resulting best-fit morphological clones are statistically similar which is expressed in overall similarity of their form (coarse-grain), but, nevertheless, difference in fine details of structural organization (fine-grain).

Here, we have shown the general logic behind the pipeline and principle possibility for generation of the morphological clones as defined above. For this purpose we used a highly variable procedural tree model (Palubicki et al., 2009), which is more difficult to optimize. As the pipeline consists of several elementary steps, each of which can be changed according to the application and target analysis, we have proposed an initial set-up and basic configuration that are capable of the task we have set. We assume larger possibilities of exploration of the proposed configuration, let alone changing the steps and individual algorithms within the pipeline, which could be fulfilled by the community of plant science researchers (for this reason, we also created a little toolbox in Matlab for easier representation and simulation of the algorithm).

Developing the principles of the pipeline, we were interested in biological plausibility of the results rather than visualization purposes. Thus, for example, we use real TLS measurements and general-purpose measure of the distance, while omitting visual effects (e.g. shades, leaves etc.). We believe this pipeline can be useful in the rigorous analysis of the plant morphogenesis and corresponding applications (in contrast to some similar studies done in computer graphics field, e.g. (Stava et al., 2014)).

Moreover, in our algorithm we employ the distance measure taking into account significant portion of the data (in fact, all data points of a given topological order(s)), not merely scalar overall entities proposed by other authors (e.g. (Frank, 2010; Stava et al., 2014)). This allows for a more comprehensive analysis of forms and their description, stemming from the statistical inference theory and in the spirit of Systems Biology studies. Due to this reason, we do not rely on the traditional metrics comparison in this work as we found that similar values for the height, girth, and crown distances may correspond to different tree forms and, thus, be non-robust.

The robustness of the statistical analysis presented here can be enhanced by using several QSM trees. In this case, similarly looking trees should be used and the degree of similarity might be established using our definition of the structural distance. For example, the trunk features are more reliably reproduced in statistical sense, when several QSM’s are used. In these lines, it might be stressed that other notions of “clone” can be used to establish relationship with morphology. Thus, the genetic clones might be utilized to establish to what degree the morphology of a tree is encoded into genes (nature vs. nurture problem).

In this initial study, we aimed at showing the plausibility of using our algorithm for effective morphology exploration. Many detailed studies scrutinizing the particulars of every part of our procedure wait to be accomplished. Among such particular questions are: QSM reconstruction configuration and its impact on the algorithm, structural distance dependence on sample size, different ways of extraction of the morphological features of a tree, multiple comparison problem, calibration of the morphological clones with QSM for the real clonal trees, use of other optimization algorithms (e.g. multi-objective ones), addressing of the “unique solution” problem etc.

## IV. Materials and methods

### Quantitative Structure Model (QSM)

QSM is derived from the point cloud obtained by TLS. Essentially, QSM is a surface reconstruction of the branches of the real tree measured by TLS. The reconstruction itself is a stochastic process, giving different architecture results for different runs. Therefore, the reconstruction introduces internal errors in addition to the TLS measurement errors. Besides giving spatial locations of parts of the tree, QSM also reconstructs topological relations between the tree branches. The branches of QSM consist of elementary units, i.e. circular cylinders, but other geometrical primitives can also be applicable (Åkerblom et al., 2015). Thus, any potential structural information about the original tree can be approximated with high accuracy with QSM (details of the reconstruction algorithm are presented in (Raumonen et al., 2013) and (Calders et al., 2015), for the validation of the algorithm see (Kaasalainen et al., 2014; Calders et al., 2015; Hackenberg et al., 2015; Raumonen et al., 2015)).

In this work, we use the reconstructed QSM of a maple tree (Fig. 2). The tree was measured in leaf-off conditions and our system consisted of a phase-based terrestrial laser scanner (Leica HDS6100 with a 650–690 nm wavelength). The distance measurement accuracy and the point separation angle of the scanner were about 2–3 mm and 0.036 degrees, respectively. The horizontal distance of the scanner to the trunk was about 7–12 m, thus the average point density on the surface of the trunk (at the level of the scanner) for a single scan is about 2–5 points per square centimeter.

The QSM of the subject maple tree consists of 19,000 cylinders approximating 3,078 branches. The tree shape was chosen due to its non-trivial form and obvious irregularities in the tree growth. This is needed to determine whether the stochastic rules of SSM growth can account for this variability (which, in fact, might come from some deterministic sources, like constant wind, shading from the neighbors, animal influences etc., and which we do not know as we do not know the history of growth). Thus, our algorithm tries to compensate the unknowns of the growth with simple stochastic rules of SSM and optimization of the stochastic distance function.

### Stochastic Structure Model (SSM)

SSM is a simulated model, preferably based on analytical and/or heuristic rules for the tree growth; however, any viable algorithm for generating tree forms may be used. Importantly, the ultimate output of the SSM simulation is a table containing data sets *U* (see IV.3 Structural data sets), describing the tree structure.

Additionally, SSM may be supplied with stochastic variability in its parameter values. Through our studies we implement simple stochastic variations (in the form of normal and uniform distributions) added to the parameter values of SSM.

Finally, the elementary units forming the SSM branches should be similar to that of QSM for the appropriate comparison or, otherwise, any differences in the form primitives must be taken into account. Usually cylinders are used in SSM studies and they were also shown, when used in QSM, to produce most reliable estimation of the real tree characteristics (Åkerblom et al., 2015).

Examples of SSM are: *LIGNUM* (Perttunen et al., 1996) – a functional-structural plant model based on the physiological principles of growth of pine trees, but also applicable to other tree forms (Lu et al., 2011); *self-organizing tree model* (Palubicki et al., 2009) is based on the heuristic principles of growth, the algorithm is capable of producing various tree shapes and is used in computer graphics; *plastic trees* (Pirk et al., 2012) are procedural growth models used in computer graphics; *AMAP/GreenLab* (see e.g. (Reffye et al., 1997; Yan et al., 2004)) is a modeling approach to generate FSPM based upon empirical rules of growth with some physiological processes taken into account.

In this work, we use self-organizing tree model (SOT) with shadow propagation algorithm (Palubicki et al., 2009) as SSM with the minimal changes as to calculate the morphological features and produce the resulting data sets for comparison with QSM (in this work we used SOT implemented in the LPFG simulator, part of the VLAB software suite, version 4.4.0-2424 for 64-bit Mac OS, see http://algorithmicbotany.org/virtual_laboratory/). This procedural tree model is fast and able to generate variety of forms, hence we can use it effectively to optimize the whole algorithm in respect to technical details as well as to cover various tree shapes. Note that more specialized tree growth models designed for the species in question would be easier subjects for the morphology optimization, but, nevertheless, can be more valuable in biologically motivated studies (the usual choice is FSPM’s, e.g. (Potapov et al., 2016)).

The total number of growth parameters of the model is 27: 23 are grouped, 4 are fixed for all times. The values of the latter are dictated both by suggestions of the authors in (Palubicki et al., 2009) and the compromise between computation time and details of the morphological description. For example, the segment length is 0.2 m (we found this optimal to grow a full size tree within a reasonable span of time, although this is not the minimum length of the target QSM segments), the voxel size is 0.2 m, and the model tree grows within 12 × 12 × 12 m cube from the center of XY plane of the cube (Z-axis is oriented upwards).

The grouped parameters are divided between 5 distinct groups corresponding to different related processes:

1. *Group I*: the initial growth parameters, including limiting values, and pipe model related parameters.
2. *Group II*: environmental effects such as sensing of the neighborhood shading, vertical gradient of the light, tropism etc.
3. *Group III*: apical dominance parameters.
4. *Group IV*: shadow propagation related constants (see (Palubicki et al., 2009)).
5. Group V: angular/branching properties.

### Structural data sets (U)

Structural data sets for any given tree structure are empirical collections of the physical dimensions and spatial orientation measures of segments and branches that are composed of segments. These data sets must be similarly obtained for any pair of {*U*_*m*_,*U*_*d*_} that is to be compared by means of the distance algorithm.

Quantities in the data sets may represent scalar characteristics and/or relations between several covariates (e.g. radii, lengths, angles, tapering function of a branch etc.). On the one hand, one needs to exhaustively describe morphology of the tree using various geometrical and topological features. On the other hand, as the number of compared data sets {*U*_*m*_,*U*_*d*_} grows the efficiency of the optimization routine decreases, since the number of distance measures to be minimized grows correspondingly (one distance value for each pair {*U*_*m*_,*U*_*d*_}). Thus, one needs more compact representation of the data. One solution is to use bigger data sets with all possibly needed (for a given application) features. (Another solution is to use multi-objective optimization routines finding, e.g. Pareto front, though we do not employ such an approach in this work.) Therefore, we use larger tables of all measured features; hence, one table represents a data set. However, we are unable to merge segment- and branch-related features into a single table as these differ in dimension (Table 1). Thus, we usually compare the array of pairs {*U*_*m*_,*U*_*d*_}, having as a result the array of distance values, but with such larger table representation we have smaller size of these arrays.

Branch– and segment-related data are described in Table 1 and Fig. 7. Throughout the manuscript we maintain the notations *B*^*w*^ and *S*^*w*^ for the branch and segment-related data sets of the (Gravelius) order *w*, respectively. The zero order *w* is assigned to the trunk (a branch connecting the tree with the ground). At the branching points, the lateral buds give rise to branches with order *w*+1, where *w* is the order of the parent branch, while the apical buds continue the branch of the same order.

**Table 1:** Branch and segment features.

| Branch features, units | Description |
| --- | --- |
| $\beta$, degree | Inclination angle of the branch, i.e. angle with its parent branch. |
| $\alpha$, degree | Azimuthal angle of the branch, i.e. angle around its parent branch (calculated from the fixed direction). |
| $L_t$, m | Total length of the branch (calculated as the sum of the segment lengths constituting the branch). |
| $R_f$, m | Initial radius of the branch, i.e. radius of its first segment. |
| $L_a$, m | Length of over the parent branch from its beginning segment to the point where the current (child) branch emanates. |
| Segment features, units | Description |
| $R$, m | Radius of the segment. |
| $L$, m | Distance from the beginning of the branch to the segment. |
| $\gamma$, degree | Angle between horizontal projections of the segment and its parent. |
| $\zeta$, degree | Angle between vertical projections of the segment and its parent. |

**Figure 7:**
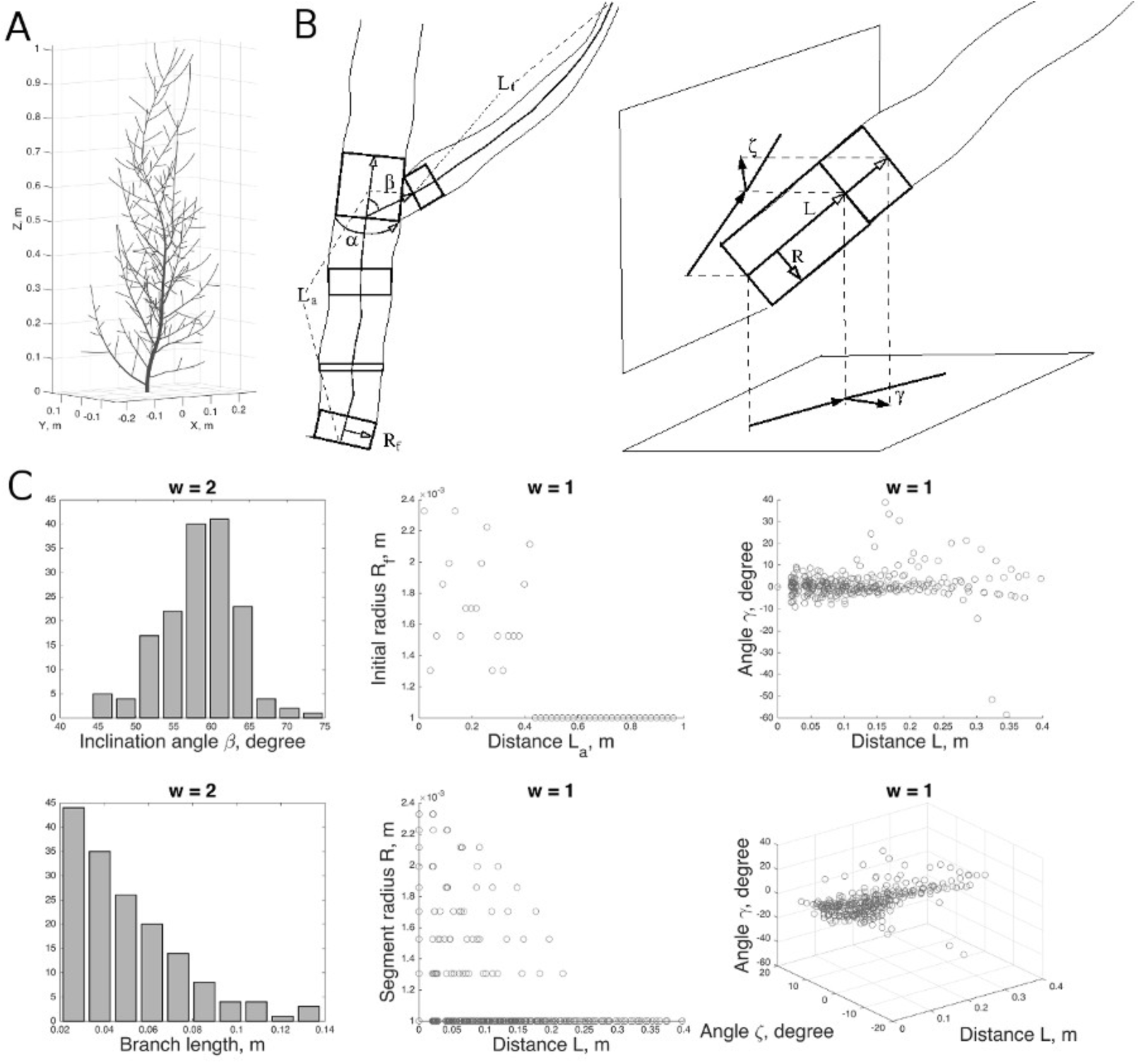
Visual structure of a tree and its representation using the structural data sets *U*. (A) A sample tree; (B) geometrical features of the branch- and segment-related data sets; and (C) various projections of the *U* data sets.

These features are not exhaustive and can be augmented at will, but we found this set sufficient for obtaining realistic tree shape outcomes. Representation of the data sets in the form of big branch and segment related tables reduces the complexity of optimization process by reducing the number of distance values to minimize. Additionally, such representation of the data allows for the fast extraction of all required relations between covariates or scalar entities without having them as separate data sets.

In a simulated SSM structure the extraction of topological relations between branches is straightforward as the user observes the whole process of growth: the lateral buds start the next order and apical buds continue the current order. However, this is not the case with QSM since it is a time snapshot of a tree form that does not retain the history of the tree growth. Thus, the reconstruction algorithm requires other principles for extraction of topology. Although the reconstruction algorithm defines a complicated procedure that outlines the topology of a tree, it could be roughly approximated by the following rule: at branching points the thickest branch is the continuation of the same order *w*, while thinner branches are lateral expansions of the order *w* + 1 (Raumonen et al., 2013). For the species with weak apical dominance (shrubby trees) we maintain similar procedure when simulating corresponding SSM (for the species with strong apical dominance, both techniques should converge to the same result).

Finally, it is possible to merge the corresponding data sets of the same order, which results at maximum in two large data sets of branch- and segment-related features, respectively. While this simplifies the search of the distance minimum (max two values to minimize), this technique must be used with care as in this case one heavily relies upon the growth rules of SSM. If these rules are not based on biologically motivated rules, SSM can produce highly unrealistic tree forms as the “best-fit”, since there is a possibility to mix the features of different topological orders. For example, the branches of higher order could be much thicker than those of the lower order, which is unrealistic and naturally is taken care of in the biologically based growth algorithms (e.g. pipe model).

### Measure of structural distance (*D*_*S*_)

The distance *D*_*S*_ between any two data sets, or empirical distributions (dimension or number of variables of which is not limited), measures the difference between the local densities of the points in *U*-space for these data sets. Here, it is constructed by measuring SSM vs. QSM difference of the normalized cumulative distributions of the point densities projected onto a number of line directions in the coordinate space of the variables in *U*. The directions of lines are generated with quasi-Monte Carlo method using low-discrepancy (quasi-/sub-random) sequences, which cover the given space more evenly than uniformly generated sequences. The difference between the projected cumulative distributions is further measured by the Kolmogorov-Smirnov statistic (any other can be used) and the resulting distance between the two data sets *U* is an average of all statistics calculated from each of the lines (see Fig. 8A).

In general,*U* ∈ *R*^*N*^, in our case *N = 4* (segment) or *N = 5* (branch) as can be seen from Table 1. The empirical probability density function *p(U)* can be approximated by the series of 1D density functions *p*_*1D*_(*U,L*), where *L* is a line in *R* ^*N*^, each of these 1D functions is constructed by projecting all the data points of *U* (thus, it is not a marginal distribution) onto a line *L* (in total we use 1000 such line directions formed quasi-randomly). Cumulative distributions *P*_*1D*_(*U*_*m*_*,L*_*i*_) and *P*_*1D*_(*U*_*d*_*,L*_*i*_) for each line direction *L*_*i*_ are compared, thus, for any given data set pair {*U*_*m*_, *U*_*d*_} the resultant distance value is:

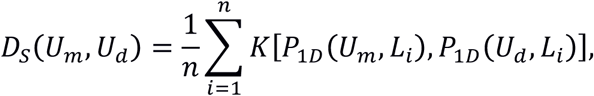

where *n* is the number of lines and operator 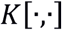 returns the Kolmogorov-Smirnov statistic for the given pair of 1D empirical cumulative distributions.

**Figure 8:**
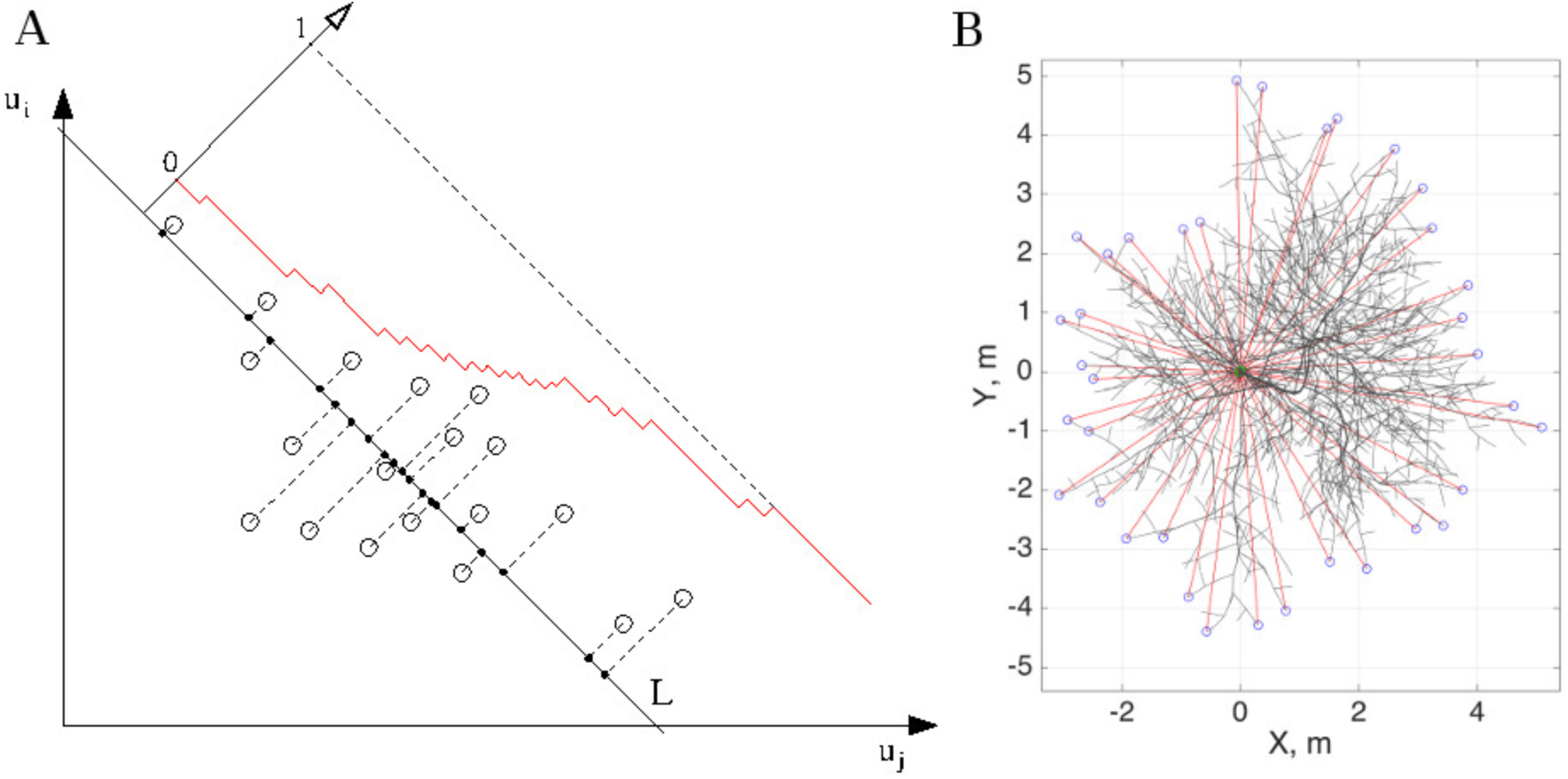
Distribution tomography of the structural data sets (A) and classical metric for the crown spread (B). (A) Data points in *U* (projected here for simplicity onto (*u*_*i*_,*u*_*j*_) plane) are used to construct the projection onto a line *L*. Cumulative empirical distribution is calculated along *L* (red). Only one line is shown, although typically one should use sufficiently enough number of lines to describe the form of the distribution. (B) Top view of a tree: spokes (red) emanate from the ground segment (green) extending up to the most distant points (blue).

**Traditional metrics (*d_x_*).** In order to provide a reference to traditional tree measurement systems, we also calculate three main tree characteristics that are used for describing a tree shape (Frank, 2010). *Height* is calculated as the highest point of a tree. *Girth* is calculated as the diameter of the ground segment (the breast-height diameter is not appropriate for the shrubby trees). *Crown spread* is calculated as follows. First, on XY-plane (top view, Fig. 8B) the set of spokes (red lines in Fig. 8B) emanating from the center of a tree (the ground segment, green circle) is formed (here, we opted for the spokes with azimuthal separation of 10 degrees). Then the length of each spoke is calculated as a distance from the tree center to the most distant point of the crown in the direction of the spoke (blue circles). The crown spread is twice an average of all spokes of a tree.

Finally, when comparing two tree shapes we calculate the distances as follows:

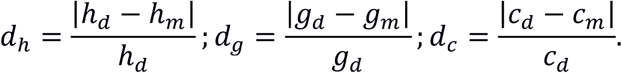

In this, *h*_*d*_, *g*_*d*_, and *c*_*d*_ are the height, girth, and crown spread of the QSM tree, respectively, whereas *h*_*m*_, *g*_*m*_, and *c*_*m*_ are the corresponding entities of the best-fit SSM tree. Thus, the classical distance *d*_*x*_ shows how large is the difference between entities *x* in proportion of the corresponding reference/QSM tree value.

### Optimization routine

The measure of structural distance *D*_*S*_(*U*_*m*_, *U*_*d*_) is minimized by adjusting the parameters *v* of SSM. In principle (with infinite sampling), *D*_*S*_ = 0 for two trees (or, more precisely, infinitely large groups of stochastically varying trees) that have exactly the same parameters *v*. These trees are not copies of each other, but they are structurally (statistically) similar. The choice of the *U* defining *D*_*S*_ is not unique, but ideally well-chosen *U* should satisfy the following uniqueness condition for *D*_*S*_ to yield an acceptable measure of distance. Let three trees be given by *v*_A_, *v*_B_, and *v*_C_. Then, if *D*_*S*_(*U*_A_,*U*_B_) < *D*_*S*_(*U*_A_,*U*_C_), one can update *C*←*B*, find any new *v*_B_ for which the inequality holds, and repeat until *D*_*S*_(*U*_A_,*U*_B_) → 0 and *v*_B_→*v*_A_. In practice, this should be true in a large enough neighborhood of *v*_A_ (any steps down the right valley lead to its bottom); however, *D*_*S*_ > 0 due to the finite sampling and insufficient model.

Any algorithm from a standard optimization library (e.g. Matlab Optimization Toolbox) that finds a minimum of an objective function (*D_S_* = *F*(*v*)) can be used. However, to facilitate global minimum search and given the nature of the problem we use the genetic algorithm (implemented in Matlab, version R2015b). Additionally, some parameters of SSM may take only integer values, so the genetic algorithm handles the integer parameters correctly unlike, for example, the classical steepest decent algorithm. The genetic algorithm iteratively finds a minimum of *D*_*S*_, each iteration being called *generation*. Each generation is characterized with a number of individuals, i.e. *population*; one individual is equivalent to one set of the parameter values. The variation is controlled by the *crossover rate* (rate of recombination of the population parameters) and *mutation rate* (rate of introduction of the new variability into the population). The former is fixed to 80% in the Matlab Optimization Toolbox, whereas the latter is controlled by our configuration. Ranges of the parameters are given by the user. There are two types of ranges: *global* lower and upper boundaries for each of the parameter values and *initial range*, from which the algorithm tries to construct the initial population (and, perhaps, where the best solution lies). The latter controls the convergence rate: if it is too broad poor convergence is attained. Finally, algorithm stops when there have passed a fixed number of generations without improving the distance.

Thus, the objective function takes the input parameters *v*, simulates SSM with *v*, calculates and returns structural data sets *U*_*m*_. Subsequently, the objective function calculates *D*_*S*_(*U*_*m*_, *U*_*d*_) and returns it to the optimization routing. The SSM, being a stochastic model, *must* have a fixed random generator seed during optimization, i.e. the same input parameter set must produce the same structural output. This is needed for convergence of the optimization. After obtaining the final best-fit form of SSM, one can further explore the variability coming from different random number sequences used in the SSM simulations (in addition to Matlab, we used GNU Octave version 4.2.0 for clone generation, see http://www.gnu.org/software/octave/doc/interpreter/). Thus, such random best-fit SSM is capable of producing the clonal morphologies (the same overall structure with varying details of organization), which is the main goal of our algorithm.

## Acknowledgments

We would like to thank Risto Sievänen and Wojtek Palubicki for useful discussion and comments on the model design and implementation.

## Competing interests

No competing interests declared.

## Author contributions

IP performed all simulations, processed the data, and wrote the manuscript; MJ wrote the code for calculating the structural distance, discussed the results; MÅ contributed to BayesForest Toolbox; PR generated and provided for the QSM data, wrote the manuscript and discussed the results; MK conceived the study, discussed the results, and wrote the manuscript.

## Funding

This work was supported by the Academy of Finland (Center of Excellence in Inverse Problem Research).

## Data availability

All data needed to reproduce the results of this study as well as some additional materials and Bayes-Forest Toolbox are available at: http://math.tut.fi/inversegroup/app/bayesforest/v1/. The most recent version of the Toolbox is also available at: http://github.com/inuritdino/BayesForest (this interface is preferred for the contributors).

## List of Symbols and Abbreviations

FSPM –: functional-structural plant model.
QSM –: quantitative structure model.
SSM –: stochastic structure model.
SOT –: self-organizing tree model.
TLS –: terrestrial laser scanning.

## References

Åkerblom, M., Raumonen, P., Kaasalainen, M., Casella, E. (2015). Analysis of Geometric Primitives in Quantitative Structure Models of Tree Stems. Remote Sensing 7(4): 4581–4603.

Bracewell, R. (1990). Numerical Transforms. Science 248: 697–704.

Calders, K., Newnham, G., Burt, A., Murphy, S., Raumonen, P., Herold, M., Culvenor, D., Avitabile, V., Disney, M., Armston, J., and Kaasalainen, M. (2015). Nondestructive estimates of above-ground biomass using terrestrial laser scanning. Methods in Ecol Evol6: 198–208.

Fourcaud, T., Zhang, X., Stokes, A., Lambers, H., and Körner, C. (2008). Plant Growth Modelling and Applications: The Increasing Importance of Plant Architecture in Growth Models. Ann Bot. 101(8): 1053–1063.

Frank, E. (2010). A Numerical Method of Plotting Tree Shapes. Bull East Nat Tree Soc6(1): 2–8.

Godin, C., Hanan, J., Kurth, W., Lacointe, A., Takenaka, A., Prusinkiewicz, P., DeJong, T., Beveridge, C., and Andrieu, B., editors (2004). Proceedings of the 4th International Workshop on Functional–Structural Plant Models, June 7-11, Montpellier, France.

Hackenberg, J., Spiecker, H., Calders, K., Disney, M., and Raumonen, P. (2015). SimpleTree - an efficient open source tool to build tree models from TLS clouds. Forests6(11): 4245–4294.

Hallé, F., Oldeman, R., and TomlinsonP. (1978). Tropical trees and forests: An architectural analysis. Berlin: Springer.

Kaasalainen, M. (2008). Dynamical Tomography of Gravitationally Bound Systems. Inverse Problems and Imaging 2: 527–546.

Kaasalainen, S., Krooks, A., Liski, J., Raumonen, P., Kaartinen, H., Kaasalainen, M., Puttonen, E., Anttila, K., and Mäkipää, R. (2014). Change Detection of Tree Biomass with Terrestrial Laser Scanning and Quantitative Structure Modeling. Remote Sensing6: 3906–3922.

LacointeA. (2000). Carbon allocation among tree organs: a review of basic processes and representation in functional–structural tree models. Ann For Sci57:521–533.

Livny, Y., Yan, F., Olson, M., Chen, B., Zhang, H., and El-Sana, J. (2010). Automatic Reconstruction of Tree Skeletal Structures from Point Clouds. ACM Transactions on Graphics 29(6): 151:1–8.

Lu, M., Nygren, P., Perttunen, J., Pallardy, S., Larsen, D. (2011). Application of the functional-structural tree model LIGNUM to growth simulation of short-rotation eastern cottonwood. Silva Fennica45(3): 431–474.

Mäkelä, A. and Hari, P. (1986). Stand growth model based on carbon uptake and allocation in individual trees. Ecol Model33: 204–229.

Palubicki, W., Horel, K., Longay, S., Runions, A., Lane, B., Mech, R., and Prusinkiewicz, P. (2009). Self-organizing tree models for image synthesis. ACM Transactions on Graphics 28(3): 58:1–10.

Perttunen, J., Sievänen, R., Nikinmaa, E., Salminen, H., Saarenmaa, H., Väkevä, J. (1996). LIGNUM: a tree model based on simple structural units. Ann Bot 77: 87–98.

Pirk, S., Stava, O., Kratt, J., Abdul Massih Said, M., Neubert, B., Mech, R., Benes, B., and Deussen, O. (2012). Plastic Trees: Interactive Self-Adapting Botanical Tree Models. ACM Transactions on Graphics 31(4): 50:1–50:10.

Potapov, I., Järvenpää, M., Åkerblom, M., Raumonen, P., and KaasalainenM. (2016). Databased stochastic modeling of tree growth and structure formation. Silva Fennica50(1), 1413.

Preuksakarn, C., Boudon, F., Ferraro, P., Durand, J.B., Nikinmaa, E., Godin, C. (2010). Reconstructing Plant Architecture from 3D Laser scanner data. In: Proceedings of the 6th International Workshop on Functional-Structural Plant Models, 14–16.

Prusinkiewicz, P. (2004). Modeling plant growth and development. Current Opinion in Plant Biology. 7(1): 79–83.

Raumonen, P., Kaasalainen, M., Åkerblom, M., Kaasalainen, S., Kaartinen, H., Vastaranta, M., Holopainen, M., Disney, M., and LewisP. (2013). Fast Automatic Precision Tree Models from Terrestrial Laser Scanner Data. Remote Sensing5: 491–520.

Raumonen, P., Casella, E., Calders, K., Murphy, S., Åkerblom, M., and Kaasalainen, M. (2015). Massive-scale Tree Modelling from TLS Data. ISPRS Annals of the Photogrammetry, Remote Sensing and Spatial Information Sciences, Volume II-3/W4, 189–196.

Rauscher, H., Isebrands, J., Host, G., Dickson, R., Dickmann, D., Crow, T., and MichaelD. (1990). ECOPHYS: An ecophysiological growth process model for juvenile poplar. Tree Physiol7: 255–281.

Reffye de, P., Fourcaud, T., Blaise, F., Barthelemy, D., Houllier, F. (1997). A functional model of tree growth and tree architecture. Silva Fennica. 1997; 31(3): 297–311.

Room, P., Hanan, J., and Prusinkiewicz, P. (1996). Virtual plants: new perspectives for ecologists, pathologists and agricultural scientists. Trends Plant Sci1:33–38.

Rosell, J., Llorens, J., Sanz, R., Arnó, J., Ribes-Dasi, M., Masip, J., Escolà, A., Camp, F., SolanellesF., Gràcia, F. et al. (2009). Obtaining the three-dimensional structure of tree orchards from remote 2D terrestrial LIDAR scanning. Agric and For Meteor 149: 1505–1515.

Rutzinger, M., Pratihast, A., Oude Elberink, S., and Vosselman, G. (2010). Detection and modelling of 3D trees from mobile laser scanning data. In: International Archives of Photogrammetry, Remote Sensing and SpatialInformation Sciences XXXVIII: 520–525.

Sachs, T. and Novoplansky, A. (1995). Tree from: architectural models do not suffice. Israel J Plant Sci43: 203–212.

Sievänen, R., Nikinmaa, E., Nygren, P., Ozier-Lafontaine, H., Perttunen, J., Hakula, H. (2000). Components of functional–structural tree models. Ann Sci57:399–412.

Sievänen, R., Perttunen, J., Nikinmaa, E., Kaitaniemi, P. (2008). Toward extension of a single tree functional-structural model of Scots pine to stand level: effect of the canopy of randomly distributed, identical trees on development of tree structure. Functional Plant Biology35: 964–975.

Smith, A., Astrup, R., Raumonen, P., Liski, J., Krooks, A., Kaasalainen, S., Åkerblom, M., and KaasalainenM. (2014). Tree Root system characterization and volume estimation by terrestrial laser scanning. Forests5(12): 3274–3294.

Stava, O., Pirk, S., Kratt, J., Chen, B., Mech, R., Deussen, O., and Benes, B. (2014). Inverse Procedural Modelling of Trees. Computer Graphics Forum33(6): 118–131.

Van Leeuwen, M. and Nieuwenhuis, M. (2010). Retrieval of forest structural parameters using lidar remote sensing. Eur J For Res129: 749–770.

Xu, H., Gossett, N., and Chen, B. (2007). Knowledge and Heuristic Based Modeling of Laser-Scanned Trees. ACM Transactions on Graphics26(4): 19.

Yan, H., Kang, M., de Reffye, P., Dingkuhn, M. (2004). A Dynamic, Architectural Plant Model Simulating Resource-dependent Growth. Annals of Botany. 2004; 93: 591–602.

